# ALSmiR: a curated evidence resource for ALS-associated microRNAs for biomarker discovery

**DOI:** 10.64898/2026.10.16.759728

**Authors:** Mariana A. Amoros, Srinivasan Ekambaram, Reda M. Iskandar, Zachary Simmons, Nikolay V. Dokholyan

## Abstract

Amyotrophic lateral sclerosis is a multifactorial, late-onset neurodegenerative disease with incompletely understood pathogenesis. Limited survival after symptom onset creates an urgent need for biomarkers to support diagnosis, prognosis, and patient stratification. MicroRNAs are among the candidate biomarkers investigated in this disease. Although reviews have synthesized microRNA expression findings, differences in patient populations, biospecimens, and methods complicate comparisons. In the age of data-driven research, we need an updated, searchable resource that links each finding to its context and incorporates growing evidence. Here, we present ALSmiR, a public, manually curated database of polymerase chain reaction-based microRNA expression findings from human biological fluids and their components. ALSmiR currently contains 1,620 expression records from 49 publications comparing patients with amyotrophic lateral sclerosis with healthy controls, covering 816 microRNA names across 13 biospecimen categories. We link publication, cohort, and microRNA-result tables to preserve each finding’s clinical and experimental context. We retain significant and non-significant results and provide searchable records and comma-separated values exports for independent analysis. We update ALSmiR annually and verify author submissions against source publications before inclusion. Researchers can use ALSmiR to assess candidate biomarkers, formulate hypotheses, and design independent validation studies. ALSmiR is freely accessible worldwide without registration at https://alsmir.dokhlab.org/.

## INTRODUCTION

Amyotrophic lateral sclerosis (ALS) is a complex late-onset neurodegenerative disease characterized by progressive dysfunction and death of upper and lower motor neurons as well as muscle weakness and atrophy^1^. ALS’s substantial clinical and molecular heterogeneity hinders efforts to understand its pathological mechanisms and develop effective therapies^2–4^. Time from symptom onset to diagnosis is estimated at 10–16 months^5^, and pooled survival time from symptom onset to death is estimated at 32.2 months^6^. This delay leaves a narrow window for disease management after a lengthy diagnosis. Clinically validated biomarkers could support earlier diagnosis, improve prognostic accuracy, support patient stratification, and increase clinical trial efficiency^7,8^.

MicroRNAs (miRNAs) are among the candidate biomarkers investigated in ALS^7^. These endogenous small non-coding RNAs regulate gene expression through translational repression and target mRNA degradation^9^. miRNAs exhibit key biomarker characteristics, including stability, non-invasive or minimally invasive sampling, and detection using accessible methods such as RT-qPCR^10^. Over the past decade, ALS miRNA research has expanded across patient populations, biospecimens, and analytical methods. Our group^11^, like others, has worked to consolidate this growing evidence^12–19^. However, these reviews cover a fixed publication period, and researchers must still consult individual studies to examine expression findings alongside their clinical and methodological context before deciding which miRNAs warrant further investigation or which gaps remain in the field.

To address this need, we developed ALSmiR, a publicly available, manually curated, and searchable database of PCR-based miRNA expression data from human biological fluids. ALSmiR complements miRBase^20^, HMDD^21^, and miRTarBase^22^, which provide miRNA sequence annotations, disease associations, and target interactions, respectively. Importantly, these resources are not ALS-specific. ALSmiR includes original peer-reviewed studies comparing ALS patients with healthy controls and adds new studies through annual literature searches and author submissions. Each curated record links a miRNA finding of increased or decreased expression, or no statistically significant difference, to its publication and to reported clinical, demographic, biospecimen, and methodological information. Researchers can assess biomarker candidates in their clinical and methodological contexts and identify understudied miRNAs, ALS populations, and biospecimens to generate new hypotheses or inform study design.

## METHODS

### Data source

We constructed ALSmiR from data extracted for our previous systematic review^11^, updated with newly eligible studies to date^13,18,23–39^. We searched PubMed, Embase, Scopus, and the Virtual Health Library through June 21, 2026, without language restrictions, using the query: (miRNAs OR microRNAs) AND (“amyotrophic lateral sclerosis” OR ALS) AND (human OR patients).

### Study selection and data extraction

We used Rayyan Essential (https://rayyan.ai) to manage literature screening, automatically detect and remove duplicates, and then activate blind mode. Two reviewers independently screened titles and abstracts using five exclusion criteria: (1) non-original studies; (2) non-ALS studies; (3) non-human studies; (4) non-miRNA expression studies; and (5) exclusively *in silico* studies. Articles that passed initial screening were assessed in full text against five eligibility criteria: (1) studies involving patients; (2) biological fluids or their components as source material; (3) comparisons between ALS patients and healthy controls; (4) PCR-based assays or validation of miRNA expression; and (5) interpretable statistical analysis. We documented the reason for each exclusion and resolved discrepancies by consensus. Eligible articles included in the previous review were excluded from the update count as “previously analyzed publication” **(Figure 1)**.

**Figure 1:**
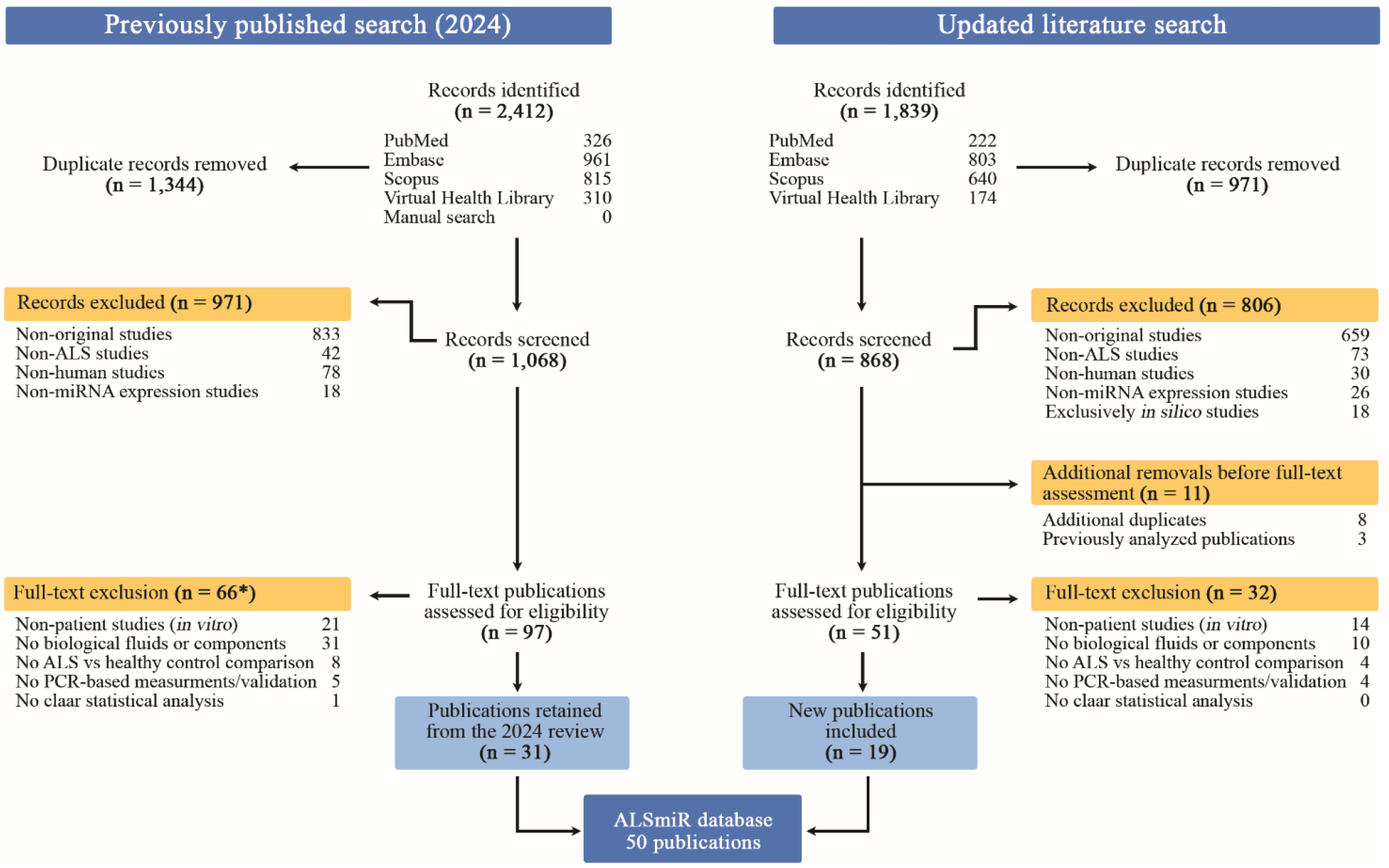
Literature search and publication screening for ALSmiR. The flow diagram shows the previously published search (left) and the updated search (right). *For the previous review, the total number of full-text exclusions is corrected from 65 to 66. ALS, amyotrophic lateral sclerosis; miRNA, microRNA; PCR, polymerase chain reaction. **Alt text:** A flow diagram shows publication selection for ALSmiR through duplicate removal, screening, and full-text assessment. The previous review contributed 31 publications, and the updated search added 18, giving a total of 49 publications.

Two reviewers independently extracted data using standardized templates, blinded to each other’s entries. We recorded publication details, clinical and demographic characteristics, and biospecimen and sampling information. We retained the authors’ reported values and units and did not infer missing information. For each miRNA result, we recorded the name and expression outcome relative to healthy controls as upregulated (↑), downregulated (↓), or not significantly different (=). We did not assign an expression outcome to missing results. When the result, figure, or text was conflicting or unclear, we contacted the authors for clarification. We recorded the number of ALS patients and healthy controls analyzed for each miRNA result, rather than assuming that the total recruited population was used. We resolved discrepancies by consensus and contacted corresponding authors when information was ambiguous or we could not reach consensus.

### Nomenclature and miRBase identifiers

A link to miRBase is provided only when the reported name exactly matches the miRBase name. Names without an exact match remain visible without a hyperlink. For selected miRNAs with ambiguous or discontinued nomenclature or missing arm information, we contacted the corresponding authors and marked confirmed revisions with “#”. We do not alter names or assign missing arms without supporting evidence. When the reported assay cannot distinguish individual miRNAs, we retain them as a group in a single entry marked with “##”.

### Web application and hosting

The ALSmiR web application is built with React (v18.3.1) and TypeScript, using Vite (v5.4.21). The interface uses Tailwind CSS (v3.4.19) and Radix UI components. Recharts (v2.15.4) and react-simple-maps (v3.0.0) provide interactive visualizations, with country boundaries from the world-atlas package. The application calculates counts and summary statistics and generates tables and visualizations client-side from the bundled database. A Node.js v20 server built with the native standard library serves the production build in a Docker container behind an nginx reverse proxy. Cloudflare provides HTTPS. ALSmiR is publicly accessible at https://alsmir.dokhlab.org.

Users can export selected evidence records as CSV files. Each exported record includes all fields associated with the selected miRNA result, its cohort, and its publication, plus two derived fields: the number of supporting evidence records and an indicator of whether the miRNA is reported in two or more publications, regardless of agreement on expression outcome.

### Database implementation

The database links three tables: publications, cohorts, and miRNA results. Each miRNA result is linked to its publication and to the specific cohort or subgroup analyzed. Cohorts reported in multiple publications appear only once and link to each publication. For each miRNA, the most frequently reported expression outcome is the category that appears in the most distinct publications. Ties indicate no unique predominant outcome. The Tissue Explorer assigns a single outcome to a miRNA-biospecimen combination when all its curated results share that outcome; combinations with multiple outcomes are classified as mixed. An independent audit compares the curated workbook with the deployed website’s records, exports, tables, summary statistics, Tissue Explorer results, biospecimen-intersection groups, and miRBase links. Automated tests compare selected website aggregations with direct calculations from the underlying dataset.

### Data submission

Researchers can submit published miRNA-ALS findings through a three-step form that collects publication and contributor information, cohort metadata, and individual miRNA results. A publication DOI is required. Each result links to a submitted cohort and specifies the expression outcome (upregulated, downregulated, or no significant change) and comparison (e.g., sporadic ALS vs. healthy controls). The form records the ALS group or subgroup, fold change, and statistical measures as reported in the source publication, without reinterpretation or normalization at submission.

The application stores submissions as structured files that preserve links among publications, cohorts, and individual results, then forwards them to curators. Curators assess eligibility and verify submitted data against the source publication before incorporating it into ALSmiR.

## RESULTS

### Database content and biospecimen coverage

As of October 2026, ALSmiR contains 1,621 miRNA expression records curated from 50 publications. These records span 816 distinct miRNA names across 13 biospecimen categories. Each record links an expression result to its source publication, cohort or subgroup characteristics, biospecimen, and PCR method.

Serum accounts for 1,246 records covering 788 miRNA names across 15 publications. Plasma contains 104 records for 45 miRNA names across nine publications, and cerebrospinal fluid contributes 66 records for 44 miRNA names across seven publications. Other biospecimen categories include peripheral blood, leukocytes, and extracellular vesicle fractions. Counts overlap when publications analyze multiple biospecimens.

### Cohort characteristics and clinical metadata availability

Of the 78 cohort records, 51 (65.4%) report fewer than 30 ALS participants, 13 (16.7%) report 30-50, and 14 (17.9%) report more than 50. Age at study is reported in 43 publications (86%), ALS functional rating scale scores in 26 (52.0%), and age at onset in 20 (40%). Eight publications record treatment information (16.0%), and six (12.0%) report the interval from disease onset to endpoint or end of life. Site-of-onset information is available for 48 of the 78 cohort records (61.5%). Bulbar onset is the most frequent category in 43 cohort records (55.1%). Availability of clinical information varies across publications, limiting interpretation of miRNA findings across patient subgroups or for stratification. To improve reporting in future ALS miRNA studies, ALSmiR recommends documenting clinical and methodological details.

### Reported miRNA expression outcomes

Of the 816 curated miRNA names, 535 (65.6%) have results from a single publication, with Waller *et al*. (2017)^40^ as the sole source for 482 of these names. The remaining 281 names (34.4%) have results from two or more publications, enabling comparisons of expression outcomes across studies.

Users can rank miRNAs by total publication count or by the number of publications reporting a selected expression outcome and display the top 6, 10, 20, or 50 miRNAs. hsa-miR-206 ranks first for increased expression in eight publications, with one additional publication reporting no dysregulation compared to healthy control **(Figure 2A)**. hsa-let-7f-5p ranks first for decreased expression in five publications, with five additional publications reporting no differences compared to control **(Figure 2B)**. Neither miRNA has reports of the opposite direction. ALSmiR retains reports for the three outcomes (upregulated, downregulated, and not dysregulated), avoiding preferential inclusion of only dysregulated miRNAs. Retaining unchanged findings allows researchers to examine where a candidate has already been investigated and consider those conditions when planning their study design.

**Figure 2:**
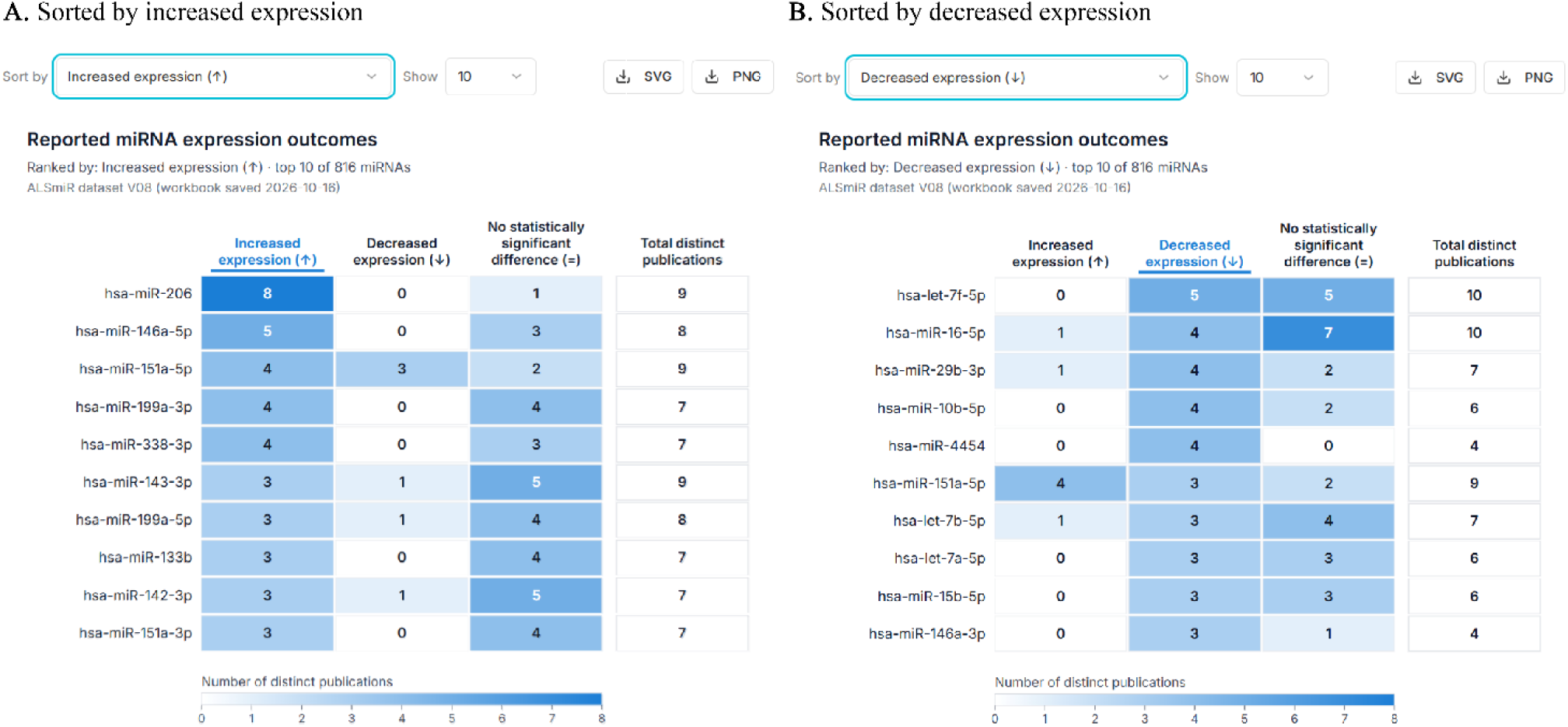
miRNAs ranked by publication counts for increased or decreased expression. The ten highest-ranked miRNAs are shown according to the number of distinct publications reporting **(A)** increased or **(B)** decreased expression relative to healthy controls. miRNA, microRNA. **Alt text:** Two heatmaps show the ten highest-ranked miRNAs by number of publications reporting increased expression (A) or decreased expression (B) relative to healthy controls. Darker blue indicates more publications. In panel A, hsa-miR-206 ranks first, with eight publications reporting increased expression. In panel B, hsa-let-7f-5p ranks first, with five publications reporting decreased expression.

### Data retrieval and exploration

A search for hsa-miR-151a-5p retrieves 15 expression records from nine publications across four biospecimen categories **(Figure 3A-B)**. The evidence table lists each comparison’s expression outcome, biospecimen, ALS cohort, PCR method, and participant counts. Expanding a record shows cohort demographics, clinical information, control-group characteristics, and publication details **(Figure 3C)**. Researchers can verify whether a finding comes from a biospecimen and patient group relevant to their own work.

**Figure 3:**
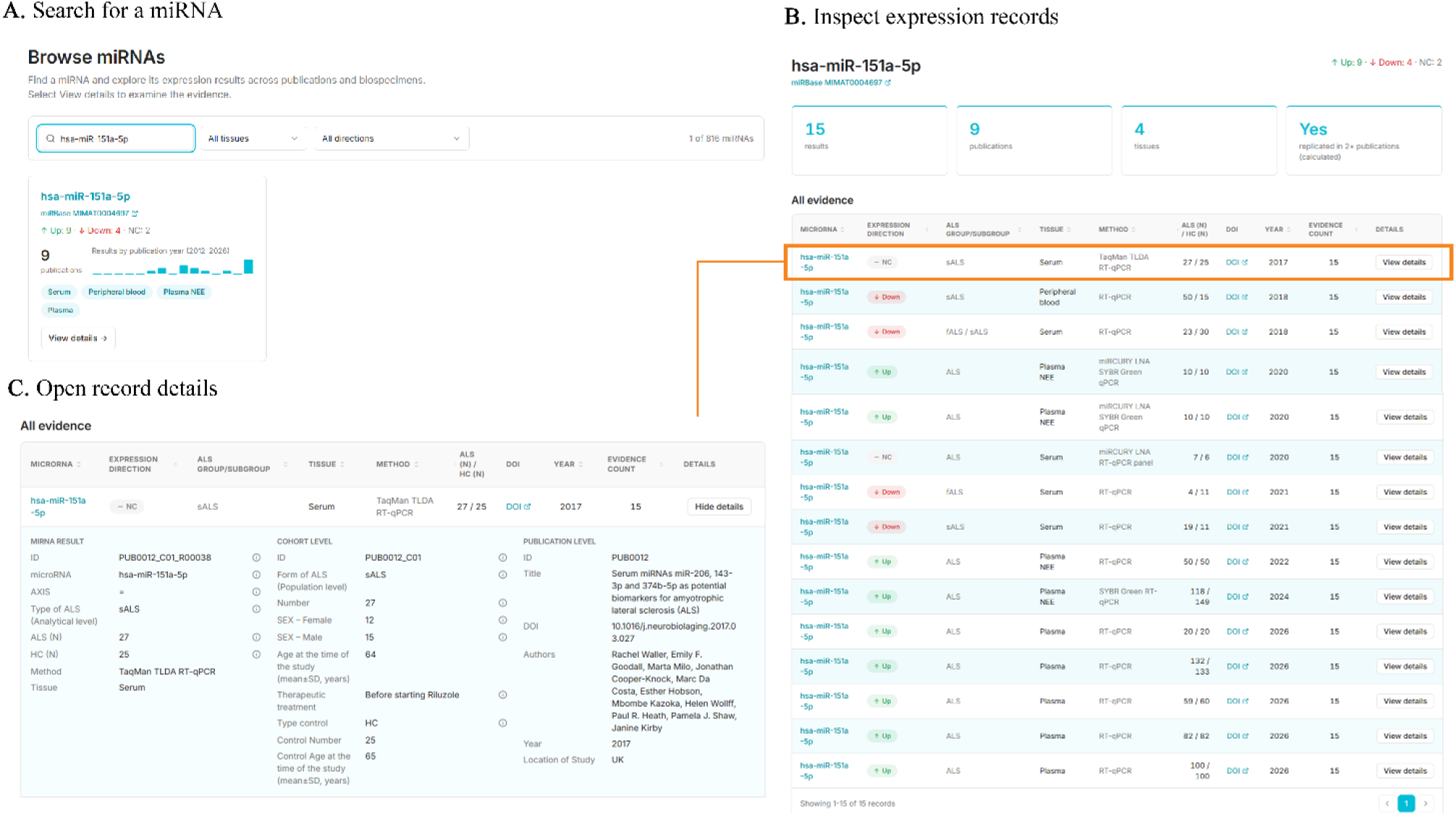
Retrieval of hsa-miR-151a-5p evidence in ALSmiR. **(A)** Search results showing the miRNA overview. **(B)** Evidence table displaying every expression record for that miRNA. **(C)** Expanded record showing study details. Expression-outcomes count refers to records, not publication number. ALS, amyotrophic lateral sclerosis; HC, healthy controls; sALS, sporadic ALS; fALS, familial ALS; NC, reported no change in expression; RT-qPCR, reverse transcription quantitative PCR; LNA, locked nucleic acid; TLDA, TaqMan low-density array; SD, standard deviation. **Alt text:** Three ALSmiR screenshots illustrate retrieval of hsa-miR-151a-5p evidence. Panel A shows the miRNA search result summary. Panel B lists 15 expression records from nine publications across four biospecimen categories: nine increased, four decreased, and two reported unchanged. Panel C shows a selected record expanded view, which displays miRNA-result, cohort, and publication details.

The Tissue Explorer separates reported outcomes for hsa-miR-151a-5p by biospecimen. All curated records for plasma and plasma neural-enriched extracellular vesicles report increased expression (Figure 4A). One peripheral blood record and three serum records report decreased expression (Figure 4B), while two serum records report unchanged expression relative to healthy controls (Figure 4C). These contrasting outcomes underscore why candidate evaluation must consider biospecimen. ALSmiR separates these findings so researchers can assess agreement within each sample type and identify miRNA-biospecimen combinations for validation.

**Figure 4:**
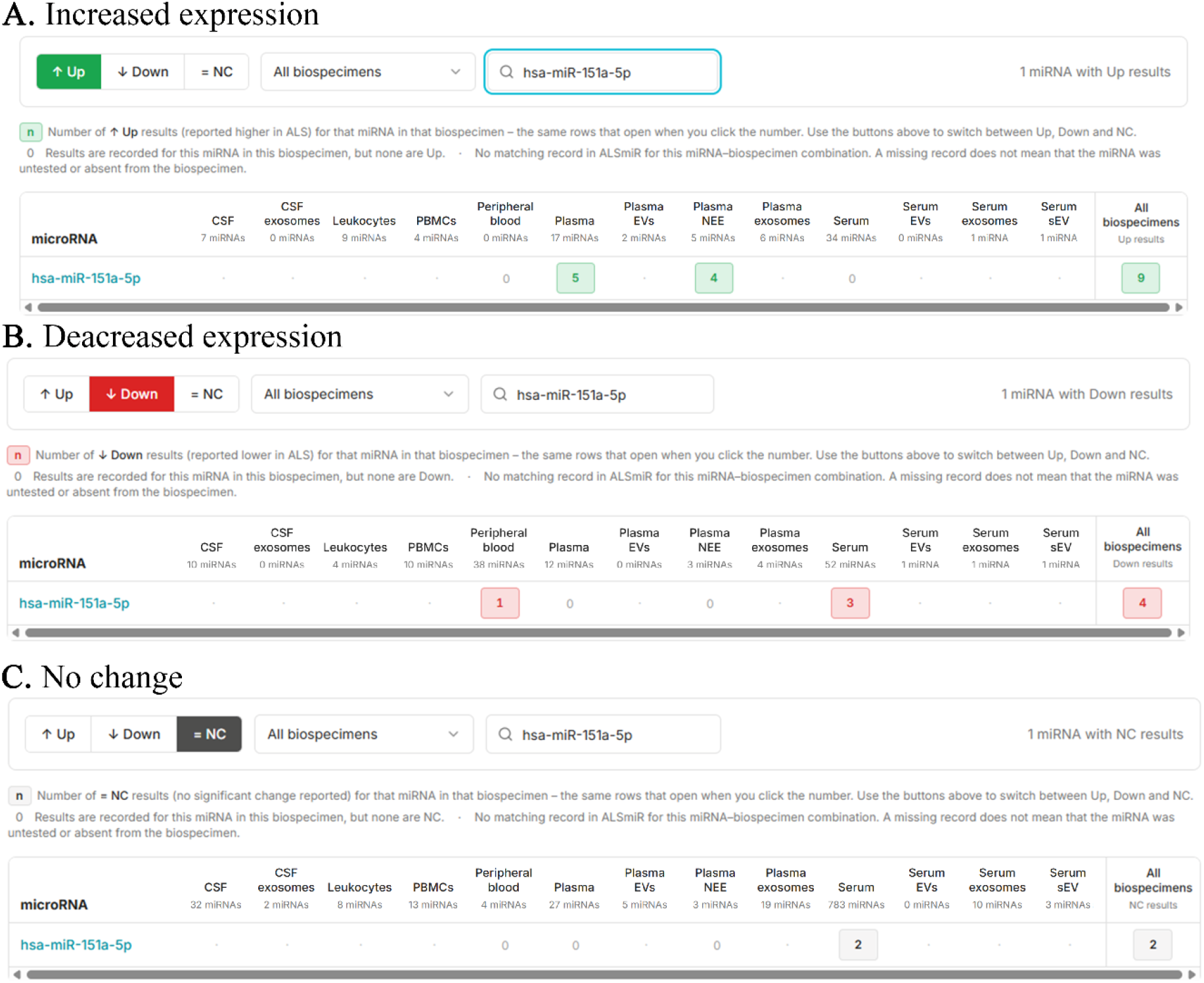
Expression outcomes for hsa-miR-151a-5p by biospecimen. Tissue Explorer views show **(A)** increased expression, **(B)** decreased expression, and **(C)** unchanged expression relative to healthy controls. Values in the miRNA row count curated expression records, not publications. A dot indicates no curated records for that combination. NC, reported unchanged expression; CSF, cerebrospinal fluid; PBMCs, peripheral blood mononuclear cells; EVs, extracellular vesicles; NEE, neural-enriched extracellular vesicles; sEV, small extracellular vesicles. **Alt text:** Three ALSmiR Tissue Explorer screenshots show hsa-miR-151a-5p expression outcomes by biospecimen. Panel A shows increased expression in five plasma records and four plasma neural-enriched extracellular vesicle records. Panel B shows decreased expression in one peripheral blood record and three serum records. Panel C shows no difference of expression compared to healthy controls in two serum records.

The four plasma neural-enriched extracellular vesicle records come from three publications by the same group^38,41,42^, whereas the five plasma records come from a single publication^35^. These patterns help researchers distinguish findings replicated across cohorts from those reproduced by independent groups and identify where further validation is needed. Users can export selected expression records, along with their cohort and publication metadata, as CSV files.

### Data integrity validation

The release audit examined all 1,620 expression records and their corresponding website outputs. The audit found no discrepancies between the curated workbook and the audited website records, exports, tables, or summaries.

## PERSPECTIVES AND CONCLUDING REMARKS

ALSmiR supports miRNA biomarker discovery by making curated evidence available to the biomedical community through a public, searchable online resource. Researchers can retrieve findings relevant to their question and export selected records, including cohort and publication information, for independent analysis. Linking each finding to its biospecimen, patient population, and methods helps researchers select candidates, design validation studies, identify gaps, and formulate new hypotheses. These links also allow users to assess the relevance of existing findings and determine where independent confirmation is needed. Conflicting findings can motivate targeted studies within a clinically defined ALS group or between tissues but should not discourage further research. Results showing no statistically significant differences require consideration of effect size and uncertainty, along with cohort characteristics and methodology.

The database supports researchers within the limits of published, peer-reviewed evidence. Missing clinical and methodological information hinders comparisons across miRNA outcomes and highlights the need for more complete reporting. ALSmiR encourages researchers to document cohort characteristics, sample collection and storage conditions, normalization procedures and statistical methods, and to report all significant and nonsignificant findings. We also encourage sharing anonymized participant-level data and numerical results in machine-readable formats to support reanalysis and future meta-analyses^11^.

Publication counts and agreement in expression direction do not establish diagnostic performance. The current focus on ALS-healthy control comparisons limits the assessment of disease specificity. We plan to address this limitation by including relevant disease controls, such as Alzheimer’s and Parkinson’s disease. We also plan to extend coverage to biopsy-derived tissues, including skeletal muscle, and to postmortem brain and spinal cord. This expansion will broaden understanding of miRNA expression patterns in ALS.

Annual literature curation and curator-reviewed community submissions will add findings to a shared, reusable record. New studies can strengthen or weaken support for an association and test whether it extends to other patient groups or biospecimens. As ALSmiR evolves, citable, archived releases are priorities to enable researchers to retrieve the database version used in an earlier analysis. These features support reproducible reuse and contribute to FAIR data practices42. ALSmiR provides the biomedical community with a shared evidence base to guide the development and independent validation of miRNA biomarkers for ALS.

## CRediT AUTHOR STATEMENT

**Mariana A. Amoros**: Investigation, Data curation, Validation, Visualization, Conceptualization, Writing – Original draft and review & editing. **Srinivasan Ekambaram:** Investigation, Conceptualization, Software, Formal analysis, Visualization, Writing – Original draft and review & editing. **Reda M. Iskandar:** Investigation, Data curation, Validation. **Zachary Simmons:** Visualization, Conceptualization, Writing – Review & editing. **Nikolay V. Dokholyan:** Visualization, Conceptualization, Supervision, Writing – Review & editing.

